# OnRamp: rapid nanopore plasmid validation

**DOI:** 10.1101/2022.03.15.484480

**Authors:** Camille Mumm, Melissa L. Drexel, Torrin L. McDonald, Adam G. Diehl, Jessica A. Switzenberg, Alan P. Boyle

**Affiliations:** Department of Human Genetics, University of Michigan, Ann Arbor, MI, USA 48109; Department of Computational Medicine and Bioinformatics, University of Michigan, Ann Arbor, MI, USA 48109

## Abstract

Recombinant plasmid vectors are versatile tools which have facilitated discoveries in molecular biology, genetics, proteomics, and many other fields. As the enzymatic and bacterial processes used to create recombinant DNA can introduce errors, sequence validation is an essential step in plasmid assembly. Sanger sequencing is the current standard for plasmid validation, however this method is limited by an inability to sequence through complex secondary structure and lacks scalability when applied to full-plasmid sequencing of multiple plasmids due to read-length limits. While next-generation sequencing (NGS) does provide full-plasmid sequencing at scale, it is impractical and costly when utilized outside of library-scale validation. Here we present OnRamp (Oxford nanopore-based Rapid Analysis of Multiplexed Plasmids), an alternative method for routine plasmid validation which combines the advantages of NGS’s full plasmid coverage and scalability with Sanger’s affordability and accessibility by leveraging nanopore’s novel long-read sequencing technology. We include customized wet-lab protocols for plasmid preparation along with a pipeline designed for analysis of read data obtained using these protocols. This analysis pipeline is built into the OnRamp webapp (http://OnRamp.BoyleLab.org), which generates alignments between actual and predicted plasmid sequences, quality scores, and read-level views in a user-friendly manner, precluding the need for programming experience in analyzing nanopore results. Here we describe the OnRamp protocols and pipeline, and demonstrate our ability to obtain full sequences from pooled plasmids while detecting sequence variation even in regions of high secondary structure, at less than half the cost of equivalent Sanger sequencing.

## Introduction

Cloning of recombinant DNA into plasmid vectors is a fundamental tool of molecular biology and central to many discoveries in genetics for decades, including the first sequencing of the human genome (Lander et al. 2001). It continues to underpin modern-day research in genomics, protein expression and purification (Rosano and Ceccarelli 2014), transcriptional regulation (Inoue and Ahituv 2015), and gene therapies (Mali 2013). However the standard for plasmid sequence validation, an important step in cloning due to the error-prone nature of recombinant assembly (Potapov and Ong 2017; Conley et al. 1986), is still Sanger sequencing, a PCR-based method invented in 1977 (Sanger et al. 1977).

Sanger sequencing uses a PCR-amplification based approach to obtain base-pair resolution of DNA sequence in stretches of up to 900bp (Sanger et al. 1977). Despite being an important tool for simple, low throughput sequence validations, Sanger also has a number of limitations. These include the need to synthesize target-specific primers, inaccuracy in long mononucleotide stretches (Shinde et al. 2003), difficulty sequencing through regions with strong secondary structure (such as repetitive elements), and a limit of about 1kb sequence output per run (Stranneheim and Lundeberg 2012). While the 900bp limit can be addressed by tiling multiple sequencing runs across the same plasmid, this requires synthesis of multiple primers and quickly becomes expensive and laborious when applied to multiple transformants. As a result, typical Sanger validation protocols involve sequencing only the portion of a plasmid that was most recently modified or that contains protein-coding sequence. However, most plasmids contain multiple elements which contribute to function (Kittleson et al. 2012); (Williams et al. 2009), including bacterial sequences (Muerdter et al. 2018), which may accumulate undetected errors as a result of spot-check sequencing approaches and can impact downstream function.

High-throughput next-generation sequencing (NGS) does allow for simultaneous sequencing of large numbers of plasmids and provides full plasmid sequences (Gallegos et al. 2020). However, NGS is cost-prohibitive outside of large-scale approaches, and sample pooling coordination, indexing compatibility issues, equipment cost, and turnaround time are major barriers to its widespread adoption and make it unsuited for routine plasmid validation. Additionally, NGS does not allow for detection of variation outside of unique regions of plasmids in libraries due to the inability to uniquely map short reads to an individual plasmid (Liu et al. 2012).

Here we present OnRamp (Oxford Nanopore-based Rapid Analysis of Multiplexed Plasmids), a tool that leverages Oxford Nanopore Technologies’ (ONT) long-read benchtop sequencing platform to obtain full sequences of pooled plasmids. OnRamp addresses the need for an approach that is simpler and more cost-effective than NGS, while providing full plasmid sequences at medium-throughput scale in a rapid, amplification- and barcode-free manner at $1.25 per kb, less than half the cost of equivalent Sanger sequencing. We describe here custom plasmid preparation protocols for use with OnRamp, testing of the OnRamp pipeline using simulated read data, and demonstrate detection of variation using real plasmid data across plasmid pools containing both dissimilar and highly similar (clonal) plasmid sequences. OnRamp comprises both these custom protocols and an analysis pipeline for ONT long-read pooled plasmid data, which is available through a web application (https://onramp.boylelab.org/), making interpretation of sequencing results accessible and simple.

## Results

### A. OnRamp protocols and pipeline

OnRamp uses ONT’s benchtop sequencing platforms that can generate reads on the order of megabases and have been employed in a variety of applications, including resolving previously intractable complex structural variation (McDonald et al. 2021), whole genome sequencing, targeted enrichment sequencing, clinical diagnostics, RNA sequencing, and metagenomics (Gilpatrick et al. 2020; Sanchis-Juan et al. 2018; Bowden et al. 2019; Wang et al. 2021). DNA sequencing using nanopore requires ligation of DNA ends with specialized adapters used to facilitate sequencing. While some have previously utilized ONT for plasmid validation, our method is unique in that it leverages full-length plasmid reads for assembly and does not require barcodes which allows for a rapid and simple sample preparation (Emiliani et al. 2022; Currin et al. 2019; Brown et al. 2022). Here we provide two methods for plasmid pool preparation for OnRamp runs based on the adapter ligation method: transposase-based or restriction digest-based **(Fig. 1A)**. The first uses ONT’s Rapid Sequencing Kit; Tn5 randomly fragments equimolar pooled plasmid DNA and simultaneously ligates ONT’s specialized sequencing adapters. In the second, plasmids are linearized with a single-cutter restriction enzyme (RE), pooled in equimolar amounts, end-repaired and mono-adenylated, then adapters are added to plasmid ends using ONT’s ligation kit. Both of these methods have different advantages. While the Tn5-based protocol is faster, the restriction-based method provides uniform, full-length DNA fragments and is used for preparation of plasmid pools containing clonal plasmid copies (see section F).

**Figure 1.**
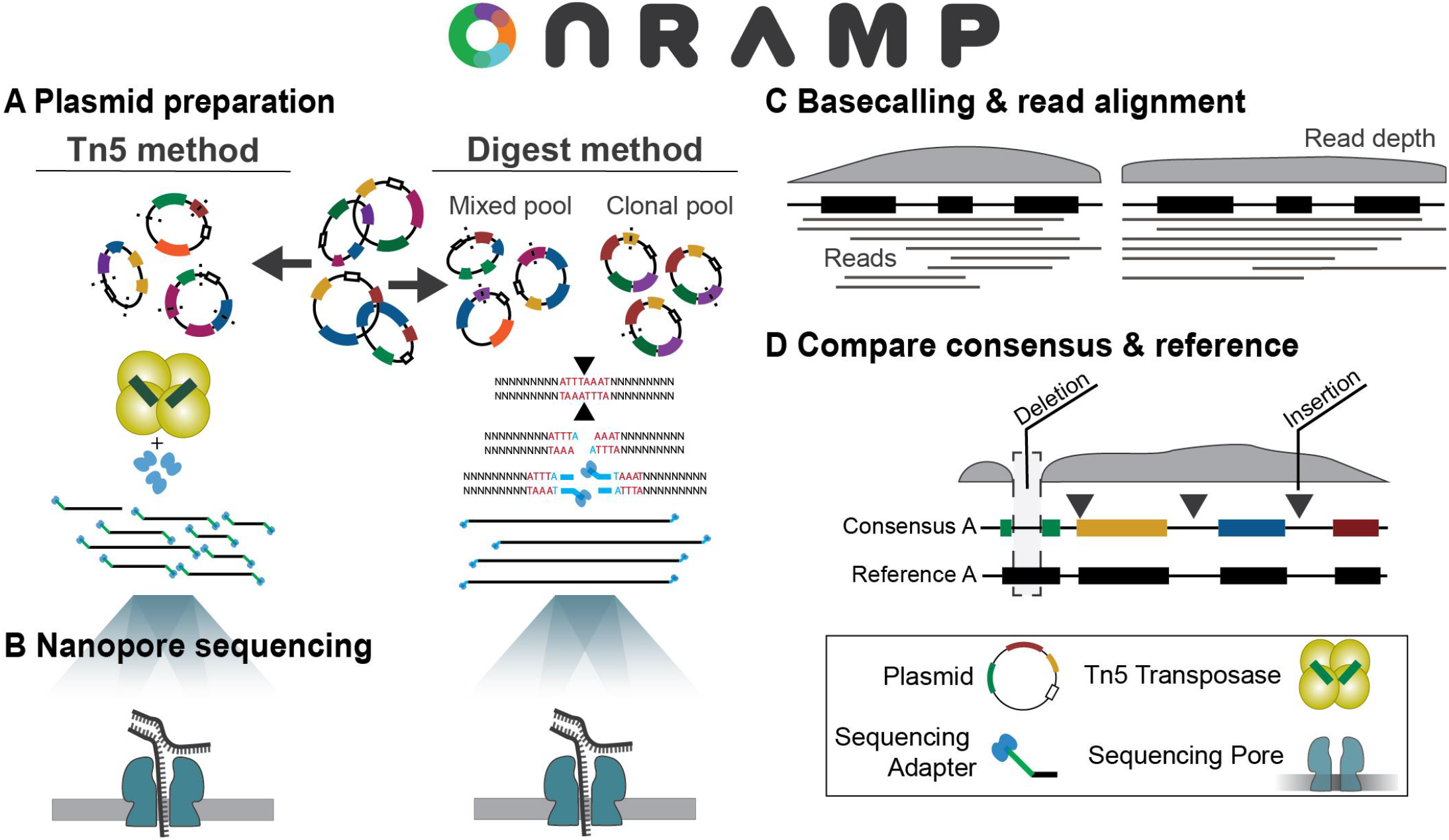
OnRamp protocol and pipeline. Pooled plasmids have Nanopore adapters added by transposase or by digestion & ligation (a) and then are sequenced (b). Basecalled reads are provided to the OnRamp webapp, which generates consensus sequences (c). Consensus sequences are then aligned to user-provided references to identify variation (d).

Following adapter ligation, plasmid libraries are loaded onto primed Flongle flow cells on ONT’s MinION platform and deeply sequenced to base-pair resolution over 16-24 hours **(Fig. 1B)** with single reads spanning entire plasmids. Basecalled read files generated by a nanopore run are then submitted to the OnRamp webapp along with plasmid reference sequence files for analysis. The pipeline run by the webapp aligns reads using medaka (https://github.com/nanoporetech/medaka) to generate a consensus sequence for each plasmid by aligning reads to user-provided references **(Fig. 1C)**, then consensuses are aligned to their matched references using Emboss Needle (Rice et al. 2000) to generate optimal global pairwise alignments **(Fig. 1D)**. After submitting a run on the OnRamp webapp **(Fig 2A)**, users are given outputs which include: a sequence-level alignment between reference and consensus files showing any insertions, deletions or base substitutions **(Fig. 2B)**, a quality score based on number and length of insertions or deletions (gaps), or base substitutions in the consensus relative to the reference **(Fig. 2C)**, and an Integrated Genome Viewer (IGV) (Thorvaldsdóttir et al. 2013) view showing read alignments used to generate the consensus **(Fig. 2D)**.

**Figure 2.**
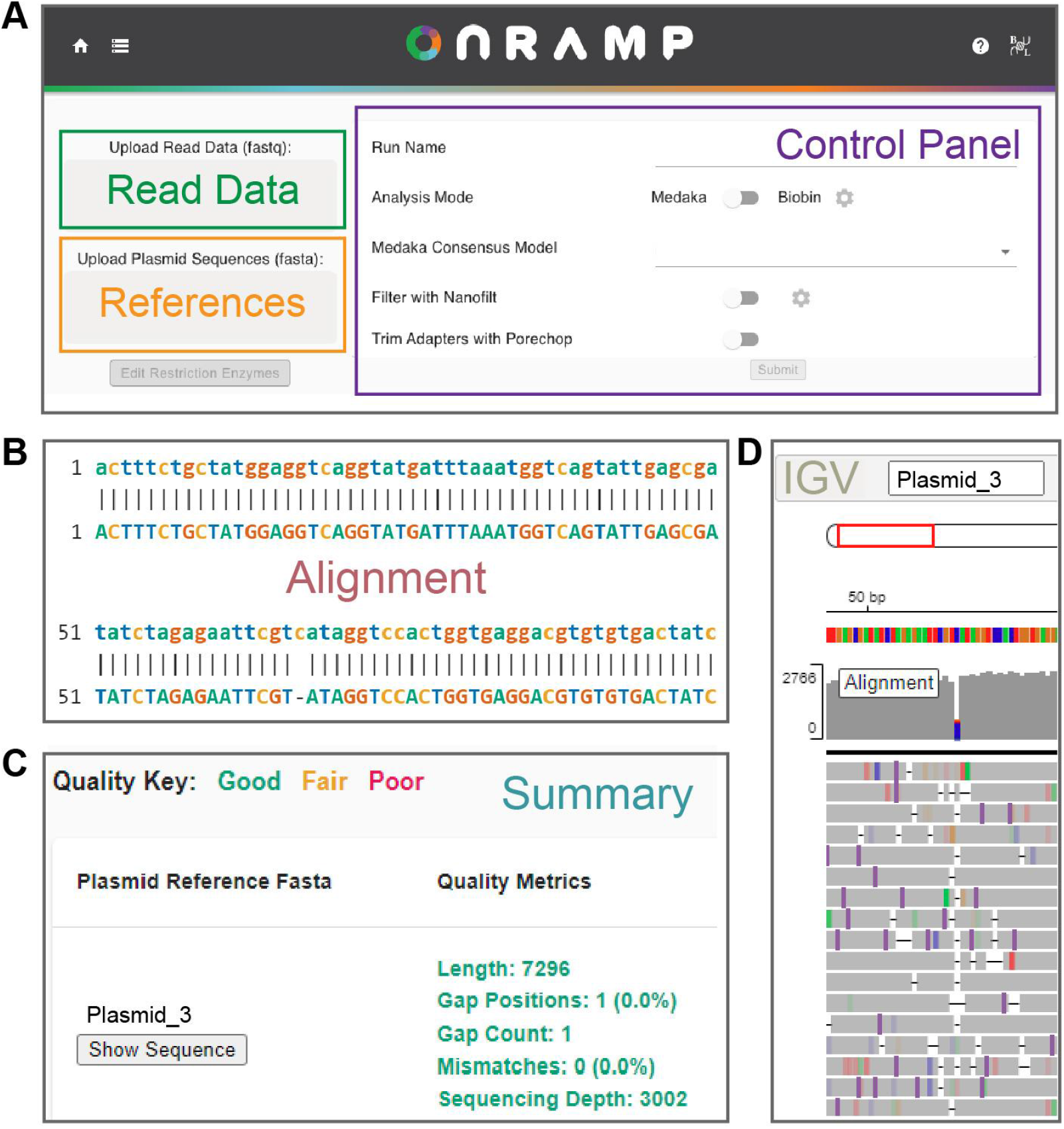
OnRamp webapp. a. Image of OnRamp submission page where users submit read data and plasmid references files and choose analysis settings. b-d. Output generated from example data, including sequence alignments (b), alignment quality metrics (c), and IGV viewer panel showing individual reads (d).

### B. OnRamp detects base-pair level variation in simulated datasets

In order to assess the ability of our OnRamp pipeline to accurately detect sequence variation occurring in plasmids from a mixed plasmid pool, we first constructed simulated read data using NanoSim (Yang et al. 2017), a tool designed to simulate nanopore reads. Read libraries were constructed for 30 dissimilar plasmids (average length 4.4kb) of known sequence, simulated to be prepared using the ONT Tn5 rapid adapter kit, giving randomly distributed read start sites. NanoSim generated 29,984 reads, with an average of 967 reads per plasmid, which were pooled and then mapped back to their respective references using OnRamp in medaka mode. In medaka mode, OnRamp uses the medaka consensus tool to generate polished consensus sequences by simultaneously mapping all reads against all references to generate best alignments, using reference sequences in place of a draft genome assembly. OnRamp mapped on average 614 reads to each plasmid, which were used to create consensus sequences. Across these 30 consensus sequences, a total of three errors (2 missing single bases at the start of one consensus due to lack of depth, and a 1bp gap at a homopolymer run in another plasmid) were observed upon alignment back to their reference sequences (1 error per 10 plasmids). Since no errors were expected given these reads originated from known sequences, we tested what level of read coverage would eliminate these gaps. We repeated consensus construction using 500%, 100%, 50% or 10% of the 29,984 reads and measured gaps in the resulting alignments **(Supp. Fig. 1A)**. Alignment accuracy varied with read coverage as expected, with more coverage giving increasing accuracy. Consensus errors consisted primarily of gaps at homopolymers and missing sequence at consensus ends due to unequal coverage across the alignments **(Supp. Fig. 3)**. Additionally, increased errors in calling homopolymers is a known limitation of ONT data (Rang et al. 2018).

Next we used this simulated dataset to test OnRamp’s ability to detect indels. A simulated read pool generated from a reference plasmid containing a 100, 10, or 1bp insertion or deletion was added to the 30-plasmid read pool. We used OnRamp to generate polished consensuses as above and results showed that insertions and deletions of 100bp, 10bp and 1bp were all correctly identified even at 100 reads per plasmid **(Supp. Fig. 1B-E)**. Read count did not impact ability to detect mutations, but rather affected whether additional variation occurred elsewhere in the consensus (points above the dotted lines in **Supp. Fig. 1D and 1E**) as a result of lack of coverage, especially at map ends and homopolymers.

### C. OnRamp correctly assigns reads to highly similar plasmids

We next used simulated data to test OnRamp’s ability to correctly assign reads originating from a pool of plasmid sequences with high sequence identity without using barcoded sequencing adapters. We created an average of 971 reads for each of 16 plasmid references differing only by 24bp, 12bp, or 6bp-long unique regions and used NanoSim to construct simulated read pools. OnRamp was then used to analyze reads and references using biobin mode. In biobin mode, OnRamp scans all provided reference sequences for unique sequence to use for distinguishing the references, then aligns each read to these unique regions to obtain an alignment score. Two tunable alignment scores, context and fine map, are used to assign each read (see methods, **Supp Fig 2A and 2B**) Each read that meets the scoring criteria is binned to this reference and then OnRamp generates a consensus for each plasmid individually from its assigned bin of reads using medaka’s consensus tool. Using biobin mode, fewer than 6% of reads were assigned to the incorrect reference **(Supp Fig 2C)** and OnRamp was able to generate consensus sequences for pools containing plasmids that differed only by the 12bp or 24bp markers. For the 6bp marker, consensus sequences contained many more gaps due to a low number of assigned reads. We also ran this test using OnRamp’s medaka mode **(Supp. Fig 2D)**, however this is not recommended as medaka mode uses non-uniquely assigned reads in consensus generation and can lead to read mis-assignment. We suggest using OnRamp’s biobin mode for correctly mapping highly similar plasmid pools where there is at least 24bp of unique sequence to differentiate the plasmids. For highly similar plasmid pools with less than ~24bp unique sequence (for example plasmids that are clonal copies), a simple alternative to the plasmid preparation protocol is provided below which works for any amount of similarity.

### D. Nanopore plasmid sequencing reveals mutations in real plasmid data

To evaluate the performance of OnRamp with sequencing of real plasmids, we ran four separate plasmid pools containing plasmids of a variety of sequences, similarity levels, and sizes, using both the Tn5 and restriction based protocols, nanopore sequenced them, and analyzed them using OnRamp’s pipeline. A 7-plasmid pool was prepared using the Tn5 transposase from ONT’s Rapid Sequencing Kit. This experiment generated 6539 reads which passed guppy’s quality filtering, an average of 934 uniquely assigned reads per plasmid **(Fig. 3A)** and a consensus accuracy of 3.4 gaps per plasmid on average **(Fig. 3B,D)**, as measured by per-base differences in consensus vs reference.

**Figure 3.**
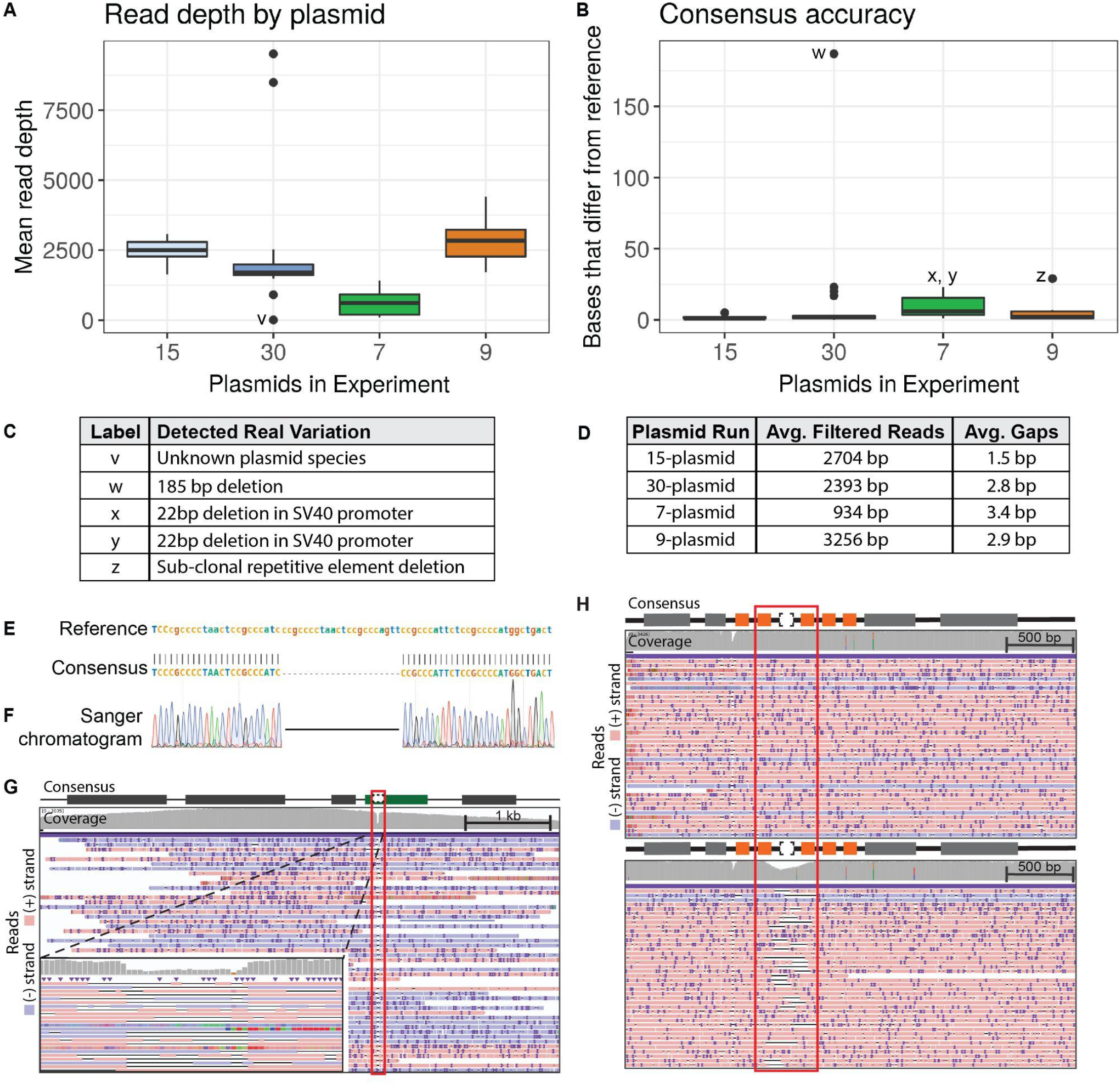
Real plasmid sequencing experiment characteristics and variant detection. a. Per-plasmid read depth across pooled sequencing runs. b. Per-plasmid count of bases in consensus sequence that differ from reference (gaps). c. Table describing variation contributing to outliers (labeled points) in a and b. d. Table summarizing read and gap data for experiments shown in a and b (gap counts do not include variants listed in c). e.OnRamp alignment results showing a 22bp deletion f. Sanger validation of deletion in (e). g. IGV browser view of reads mapping to deletion (red outline) from (e) in an SV40 promoter (green box), Left inset: zoomed view. h. IGV view of reads mapping to a clone without (top) or a clone with (bottom) a sub-clonal repetitive element (orange boxes) deletion (red outline). IGV: Black lines are deletions, dark purple marks are insertions.

The high read coverage and read length generated allowed us to distinguish reads and generate consensuses from three highly similar plasmids in this run that differed only by two 4bp sequences. Additionally, real sequence variation was detected in this run (not included in the per-plasmid gap average). A 22bp deletion, too small for detection by diagnostic digest & gel electrophoresis, was identified in the SV40 promoter of two plasmids **(Fig. 3C,E)** and validated by Sanger **(Fig. 3E-G)**. This deletion occurred outside of a region manipulated by molecular cloning and would not normally have been checked and caught by Sanger sequencing.

The 9-, 15-, and 30-plasmid pools were prepared using the restriction digest & ligation method. In the 9-plasmid pool, plasmids had over 1000 reads per plasmid, with on average 3256 quality filtered reads per plasmid, and an average of 2.9 gaps per plasmid consensus **(Fig. 3A-D)**. In this run, we were able to sequence through 4× and 6× 40bp repeats in 6 of the plasmids that were previously intractable to Sanger sequencing due to high secondary structure. The high read coverage obtained on these runs allows us to identify sub-clonal populations in plasmid sequences, which can occur as a result of plasmid recombination during bacterial growth. We detected a sub-population level deletion of one of these repeats in a plasmid from this run using OnRamp **(Fig. 3C,H)**. This high coverage is reflected even in the 30-plasmid pool, which averaged 2393 quality filtered reads per plasmid, and minimum of 900 reads per plasmid, generating base-pair resolution for all but one plasmid. This single exception was expected as it was known to have failed a diagnostic restriction digest check, indicating its sequence likely would not match the provided reference. This run had 2.8 gaps per plasmid on average, excluding a 185bp deletion detected in one plasmid **(Fig. 3B,C)**.

### E. Validating plasmid sequences in pooled plasmid clones

The 9- and 15-plasmid runs described above all contained some plasmids that were clonal copies of each other. Normally, as a result of plasmid pooling, reads originating from different clones of the same plasmid (or highly similar plasmids) would all map to the same reference, making differentiation of the read source impossible without barcoding. However, we were able to successfully leverage nanopore’s long read length in a simple alternative restriction-based protocol to differentiate reads originating from identical clones in the same pool without the need for barcoding. For plasmid libraries containing multiple plasmid clones or highly similar plasmids (<=24 bp difference), each clone is cut with a different unique restriction enzyme from its matched partners prior to pooling **(Fig. 4A)**. During analysis, a copy of the same plasmid reference sequence is provided for each clone, except with the linear sequence origin set at the digest site used for that clone (termed ‘rotated’ reference). While each cut clone contains the same total sequence, the alternate digest sites create linear fragments (reads) that map precisely to their matched ‘cut’ reference sequence, but poorly to the same sequence reference ‘cut’ at any other site (**Fig. 4A**). This approach is feasible due to the long-read nature of nanopore sequencing, where the majority of reads span an entire plasmid.

**Figure 4.**
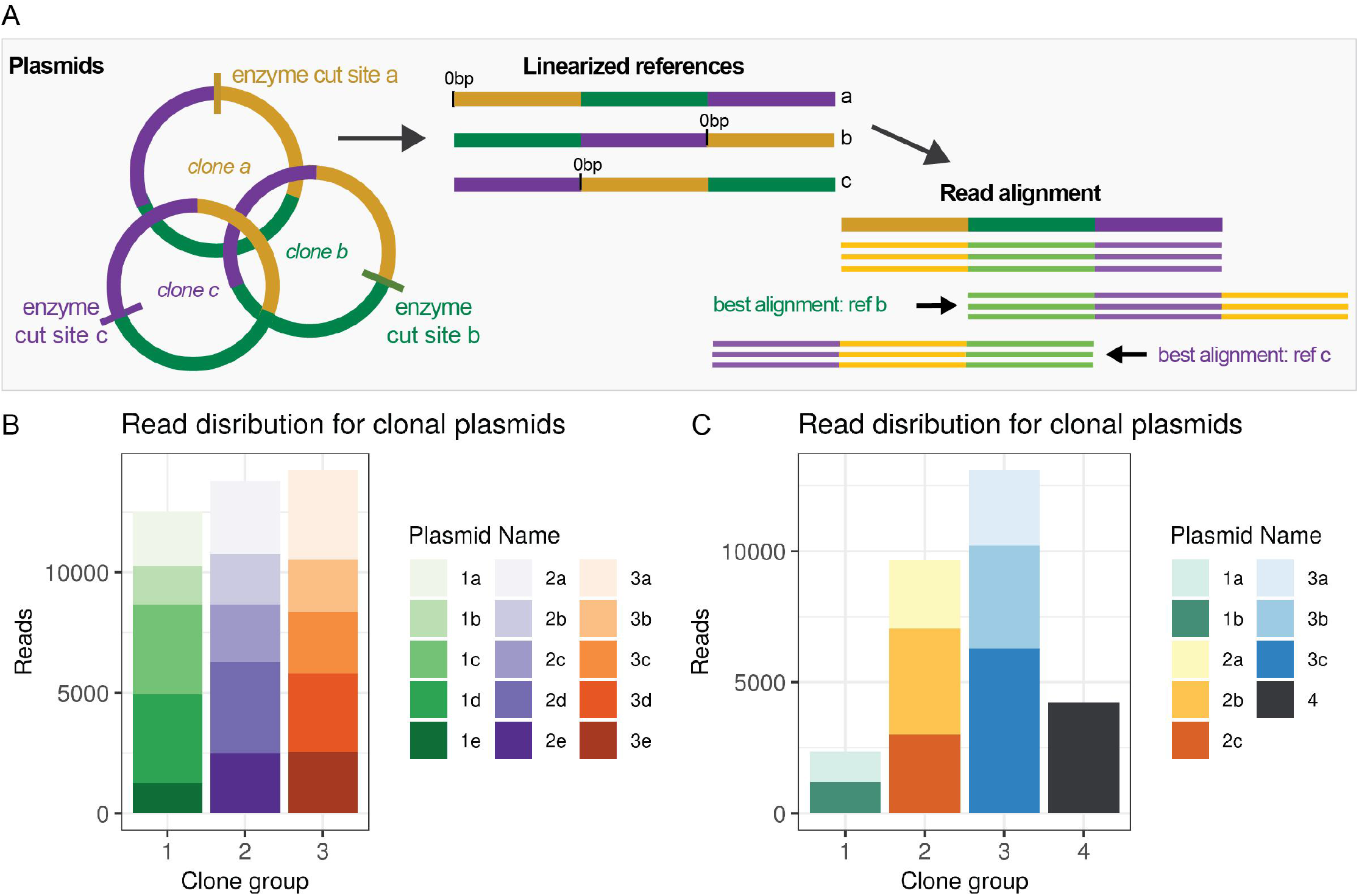
Restriction-digest barcoding for highly similar or clonal plasmids. a. Diagram of restriction cut-site method for unique read mapping of clonal plasmids using 'rotated ' references. b. Number of reads mapping uniquely to each plasmid in a 15-plasmid clonal test pool. c. Reads mapping uniquely to each reference in a 9-plasmid mixed clonal run.

We validated this approach experimentally using three ~6.5kb plasmids **(1,2 and 3 in Fig. 4B)** which are identical except for a ~500bp region. Five clones of each plasmid **(a-e in Fig. 4B)** were digested using different restriction enzymes for clones within a set, with the closest cut sites 579bp apart. An average of 2704 reads uniquely mapped when using rotated references **(Fig. 4B)**, compared to 7 reads uniquely mapping to non-rotated references, indicating that using different cut locations with clones is sufficient to create reads that align uniquely to their matched rotated reference. The 9-plasmid run contained 3 sets of clones and one unique plasmid **(Fig. 4C)**. The sub-population level deletion of a repetitive element discussed above **(Fig. 3G)** was detected in a plasmid that was part of a set of three clones in this run.

## Discussion

Assessing recombinant plasmid sequence fidelity is an integral part of any molecular cloning workflow. While Sanger sequencing is an elegant, cost-effective method for low-throughput plasmid validation, it can be inadequate for multi- whole-plasmid sequencing, and handles regions with complex secondary structure poorly. As an alternative, high-throughput next-generation sequencing’s run cost, equipment cost and sample coordination requirements make it inefficient for standard plasmid validation workflows outside of large plasmid libraries. Additionally, NGS requires amplification and due to its short-read nature, cannot identify and correctly assign mutations outside unique regions in highly similar plasmid pools. With the introduction of Oxford Nanopore Technologies’ sequencing platforms, sequencing of many plasmids in their entirety at high read depth is now possible. While some techniques have been published for recombinant plasmid verification using ONT, they rely on Tn5 barcoded libraries and *de novo* assembly to validate plasmid libraries (Emiliani et al. 2022; Currin et al. 2019).

OnRamp combines both wet-lab protocols for pooled plasmid preparation using the ONT platform, and an associated reference-based computational pipeline packaged in the OnRamp webapp, which is designed specifically to support validation of plasmid pools. ONT’s compact benchtop sequencing platform is much more affordable than most NGS sequencing platforms and allows for in-lab sequencing with results available as soon as next-day, without the need to ship samples to a core or company. OnRamp provides a rapid (0.5-2 hours for preparation, 16-48 hours for results) and cost-effective approach for medium-throughput plasmid sequencing. Using the reagents and protocols described here, we were able to fully sequence 30 plasmids at $1.25 per kb, significantly less than the cost of using Sanger sequencing to obtain equivalent data. Additionally, the OnRamp webapp facilitates analysis and data interpretation in a manner that is accessible to labs without the need for extensive bioinformatics support.

Testing OnRamp using simulated read libraries demonstrated its ability to correctly assign sequencing reads to reference sequences and construct consensus sequences even with highly similar plasmids. Testing OnRamp on real plasmid runs showed that OnRamp provides high sequence read depth across plasmid pools, generating consensus sequences spanning entire plasmid lengths at base pair resolution (even on Flongle flow cells with as few as 20 pores). The read depth we obtained (over 900 reads per plasmid for a 30-plasmid pool) using even ONT’s lowest-capacity flow cell (the Flongle), allows for high confidence in base-level calls in consensus sequences despite nanopore’s 10% error rate (Chandak et al. 2020). This makes the level of difficulty of interpretation of OnRamp results about the same as that of Sanger sequencing. OnRamp’s output includes IGV viewer alignments of individual read data for interrogating the underlying data to determine confidence in consensus sequence base calls. The high coverage we obtained allows for detection of sub-population level variation even in a region with complex secondary structure and high clonal similarity. Additionally, this mutation (a deletion of one of a 6x repetitive element set) likely occurred as a result of bacterial recombination that occurred after cloning, underscoring the importance of obtaining full-plasmid sequencing data rather than running spot-check validations using Sanger. These experiments also revealed a 22bp mutation in a functional non-coding plasmid element (SV40 promoter) which was previously undetected by diagnostic restriction digests, showcasing the ability of the tool to determine uncharacterized structural and sequence variation.

Using OnRamp’s medaka mode to generate consensus sequences, we were able to rapidly validate our plasmids based on alignments to reference sequences. Some limitations of this approach arise as a result of medaka being a reference-based approach, as opposed to an assembly-based method. OnRamp uses a reference-based system for analysis, as it is a tool designed specifically for routine plasmid validation. While this precludes use of OnRamp for *de novo* assembly of unknown plasmid sequences (an uncommon case in routine screening), there already exist well-designed tools for *de novo* assembly of nanopore data (Emiliani et al. 2022; Currin et al. 2019; Brown et al. 2022). For instance, while we were able to detect most variation in our constructs, consensus sequences for plasmids with very large indels (>1000bp) or where large portions of the plasmid have inserted backwards relative to the reference could not be generated. However, these large rearrangements should be easily detected by complementary diagnostic restriction digest tests, which are often a routine step in cloning protocols. Using the alternate biobin mode, choosing unique regions in the reference is essential to binning reads. Indels in the unique portion of the reference can lead to incorrectly binned reads or failure to generate a consensus. An alternative method is to use the clonal restriction-based method we described to separate reads from highly similar plasmids.

We found instances of mixed plasmid populations post-transformation **(Fig. 3D)**. This mixed population can be missed in the consensus file however, this issue can be addressed by viewing the read alignments in IGV (provided as part of the output for any run using the OnRamp webapp) or a similar program. This problem also appears in Sanger sequencing results where sequence files will not show sub-population structure, but could be detectable in trace files. OnRamp offers rapid, medium-throughput full-plasmid sequencing without secondary structure limitations or the need for primers. It provides more affordability and simplicity than NGS, and with our streamlined web application, makes analysis and interpretation of results accessible and straightforward.

## Methods

### Vector construction and maintenance

Plasmids were constructed using either EMMA (Martella et al. 2017) or gateway- or restriction-based cloning methods. The EMMA toolkit was a gift from Yizhi Cai (Addgene kit # 1000000119). Various parts from the toolkit were used for construction of the vectors, and mCherry was cloned from pHR-SFFV-KRAB-dCas9-P2A-mCherry to become a usable part. pHR-SFFV-KRAB-dCas9-P2A-mCherry was a gift from Jonathan Weissman (Addgene plasmid # 60954; http://n2t.net/addgene:60954; RRID:Addgene_60954) (Gilbert et al. 2014). Expression vectors were grown in either Stbl3 or DH5ɑ chemically competent E. coli strains.

### Tn5-based plasmid preparation

For TN5-based preparation, plasmids were treated using the Rapid sequencing kit and following ONT’s protocol (ONT, SQK-RAD004). Pooled plasmid DNA is brought to 7.5uL using H2O and combined with 2.5uL FRA, and incubated 30°C for 1 minute and then at 80°C for 1 minute then put on ice. 1uL of RAP is added and mixed by flicking, then spun down and incubated for 5 minutes at room temperature. DNA is loaded onto a primed flow cell.

### Plasmid pool linearization by restriction enzyme and end-repair

Plasmid DNA was isolated using the QIAprep Spin Miniprep Kit following the manufacturer’s protocol (QIAGEN, 27104) and eluted in water. Plasmids were linearized by restriction digest using a unique cut site, with times, temperatures and reaction volumes varied for other enzymes according to NEB recommendations. Example pooled restriction digest: NEB Buffer 3.1 (NEB,B7203S) was added to 1X and the final volume was adjusted with nuclease free water to 200uL. SwaI (NEB, R0604L) was added according to the total amount of DNA present for linearization (minimum 10 Units enzyme per 1 ug DNA), and the sample was digested at 25°C for 30 minutes. Plasmid pools were generated prior to digest if all contained the same unique restriction site, or after digest for plasmid pools where each plasmid required a different restriction enzyme. For plasmids where different restriction enzymes are used on each plasmid, heat-inactivation of each enzyme (following manufacturer instructions) or if not possible, column cleanup (Qiaquick PCR purification kit, QIAGEN, 28104) to remove enzyme was done and is a crucial step prior to pooling to prevent cross-cutting of other plasmids in the pool after combination by still-active enzymes.

Digested plasmids were diluted and pooled into a single 1.5mL microcentrifuge tube using the following rules to calculate desired amount of each plasmid: 1. using an equimolar amount of each plasmid, 2. a maximum of 1000ng total plasmid for the entire pool 3. using at least 10ng of each plasmid 4. a total 50uL volume. Amount of each plasmid in a pool ranged from 15ng-100ng across experiments in this paper. If any digests generated 3’ or 5’ overhanging bases, pooled plasmids were end-repaired using 1uL (5U) DNA Polymerase I Klenow Fragment (NEB M0210S) with 33 μM each dNTP and 1x NEB CutSmart buffer per 1000ng DNA pool, with incubation for 15 minutes at 25°C, and heat inactivation for 20 minutes at 75°C. Following digestion and end repair, A-tailing was completed using 1uL of 10mM dATP and Taq DNA polymerase (NEB, M0273S) per 50uL of sample with incubation at 75°C for 15 minutes.

### ONT Adaptor Ligation

For restriction-prepared enzymes, following DNA linearization, end-repair and A-tailing, ONT’s ligation sequencing kit was used (ONT, SQK-LSK109) to add adaptors. One half volume of ligation buffer (4X T4 ligase buffer, 60% PEG 8000), 5uL of T4 DNA ligase (NEB, M0202M), and 2.5uL of AMX (ONT, SQK-LSK109) was added to the plasmid mixture then incubated on a tube rotator at room temperature for 10 minutes. One volume of 1X Tris-EDTA buffer (pH 7.5; Invitrogen, 15567027) and 0.3X room temperature SPRI beads (Beckman Coulter, B23317) were added for selection of >2 kb fragments. The sample-SPRI bead mix was incubated on a tube rotator for 10 minutes on the bench at room temperature. The SPRI beads were washed twice with 100uL of Long Fragment Buffer (LFB; ONT, SQK-LSK109) and the sample was eluted in 9uL of Elution Buffer (EB; ONT, SQK-LSK109).

### Nanopore sequencing

Flongle flow cells were primed for the sequencing runs following ONT’s standard protocol, using flow cell priming buffers provided by ONT. Briefly, flow cells are QCd to check for a usable number of active pores (~ 0.5-1 pores per plasmid was used here as the minimum). Flow cell was washed with FB then SQB buffer mixed 1:1 with water. DNA prepared from previous steps is mixed with SQB and LB immediately prior to loading following ONT’s protocols.

### Simulated reads

NanoSim was used to construct pooled plasmid read libraries. First, a model was created using 81,070 reads (N50=6,003bp) from a previous plasmid sequencing experiment, and the 30 plasmid sequences (average length = 4,318.7bp) were used as the reference genome and input in the characterization set. This model was then used to simulate reads from other plasmid references and from references constructed with 1, 10, 100, and 1000bp deletions and insertions of random sequence as well as plasmids with 6, 12, 24bp unique regions.

### Bioinformatics pipeline

Basecalling was completed using Guppy (Oxford Nanopore Technologies, 4.5.2) using the dna_r9.4.1_450bps_hac.cfg configuration and passing reads (Q >= 7) were filtered using Guppy or NanoFilt (De Coster et al. 2018). Adapters were trimmed using Porechop (https://github.com/rrwick/Porechop). OnRamp allows users to use Porechop and NanoFilt to trim reads and filter by q score and read length. Reference sequences were generated using SnapGene (https://www.snapgene.com/). The reads and references were then used as input for OnRamp during testing.

### Medaka

The medaka consensus (https://github.com/nanoporetech/medaka) module was utilized to generate consensus sequences from read pileups using the ‘-g’ flag to stop filling in gaps with draft/reference sequence during consensus stitching. The resulting alignments are then filtered (MAPQ >= 10) for visualization using the Integrative Genomics Viewer (IGV) (Thorvaldsdóttir et al. 2013). Final pairwise alignments were constructed between the reference and consensus sequences generated by medaka using Emboss needle.

### Binning

The biobin module mode of plasmid sequencing was used to bin reads based on unique sequences in the provided references. The biobin mode/module searches the reference sequences for unique sequences longer than 3bp and a set is constructed for each plasmid reference. Each input read was then aligned to these regions using Biopython pairwise aligner with alignment parameters match: 3, mismatch: −6, open_gap: −10, extend: −5. Reads were first aligned to an extended portion of the plasmid containing 20bp flanking the unique region and assessed using the ‘context score’. For reads that passed this threshold, the aligned portion was then aligned and scored against the exact unique region and high scoring reads (fine score > 80) were assigned to the plasmids. Each of the resulting bins was then passed to medaka for consensus polishing.

## Data Access

The OnRamp is available through a webapp at https://onramp.boylelab.org/. The command line version and pipeline used for the application are available at https://github.com/Boyle-Lab/bulk_plasmid_seq_web and https://github.com/crmumm/bulkPlasmidSeq.

## Acknowledgements

M.D. and T.M. were supported by NIH Training Grant Michigan Predoctoral Training in Genetics (T32GM007544) C.M was supported by University of Michigan Genome Science Training Program (T32HG000040).

## Author Contributions

A.P.B., T.M., M.D. and C.M conceived the project. C.M and A.G.D. developed OnRamp pipeline and webapp. C.M performed data analysis. All authors guided the experiment design and data analysis strategy (C.M conceived and performed simulated experiments, T.M conceived and performed Tn5 in-vitro experiments, M.D. conceived and performed in-vitro clonal plasmid sequencing experiment, and J.S and M.D ran the other restriction-based in-vitro sequencing experiments). A.P.B. supervised the experiments, analysis, and data interpretation. C.M, M.D. and T.M wrote the manuscript and all authors contributed edits and revisions. All authors read and approved the final manuscript.

## Supplement for “OnRamp: rapid nanopore plasmid validation”

**Supplemental Figure 1.**
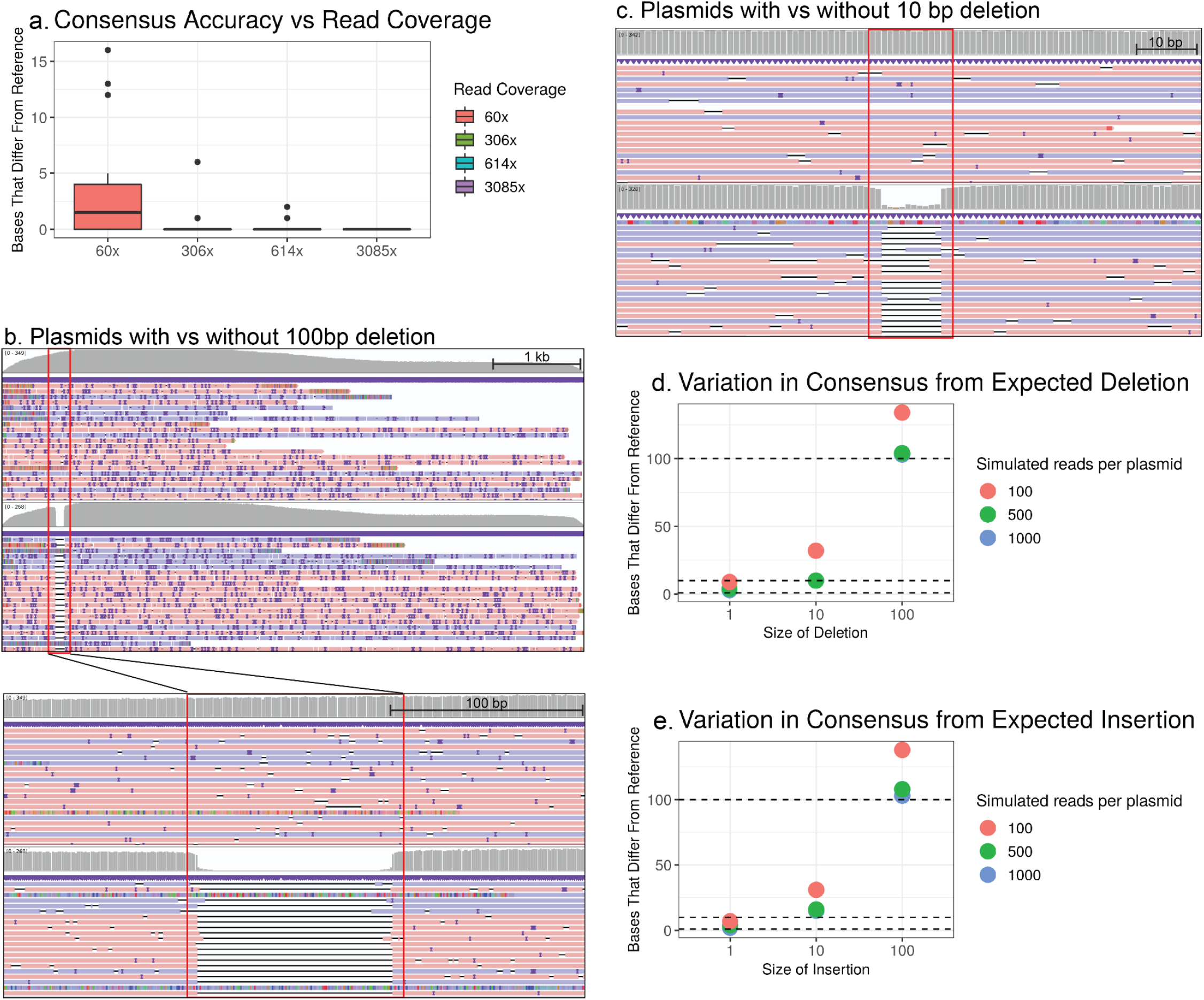
Detecting insertions and deletions in a plasmid pool using a simulated read library. a. Number of gaps in consensus sequence (before indels) for simulated read depth experiments at various read coverage. b and c. IGV view of read pileups for reads with vs reads without a 100 bp deletion (b) and a 10 bp deletion (c). Deletions are highlighted by red boxes. Gray top row shows read depth at each position. Purple lines are minus-strand reads, red lines represent plus-strand reads. Dark purple = inserts, black lines = deletions. d and e. Number of base-pair differences between reference and consensus files for each simulation condition at different read depths. Dotted lines indicate expected number of differences due to simulated deletion (d) or insertion (e).

**Supplemental Figure 2.**
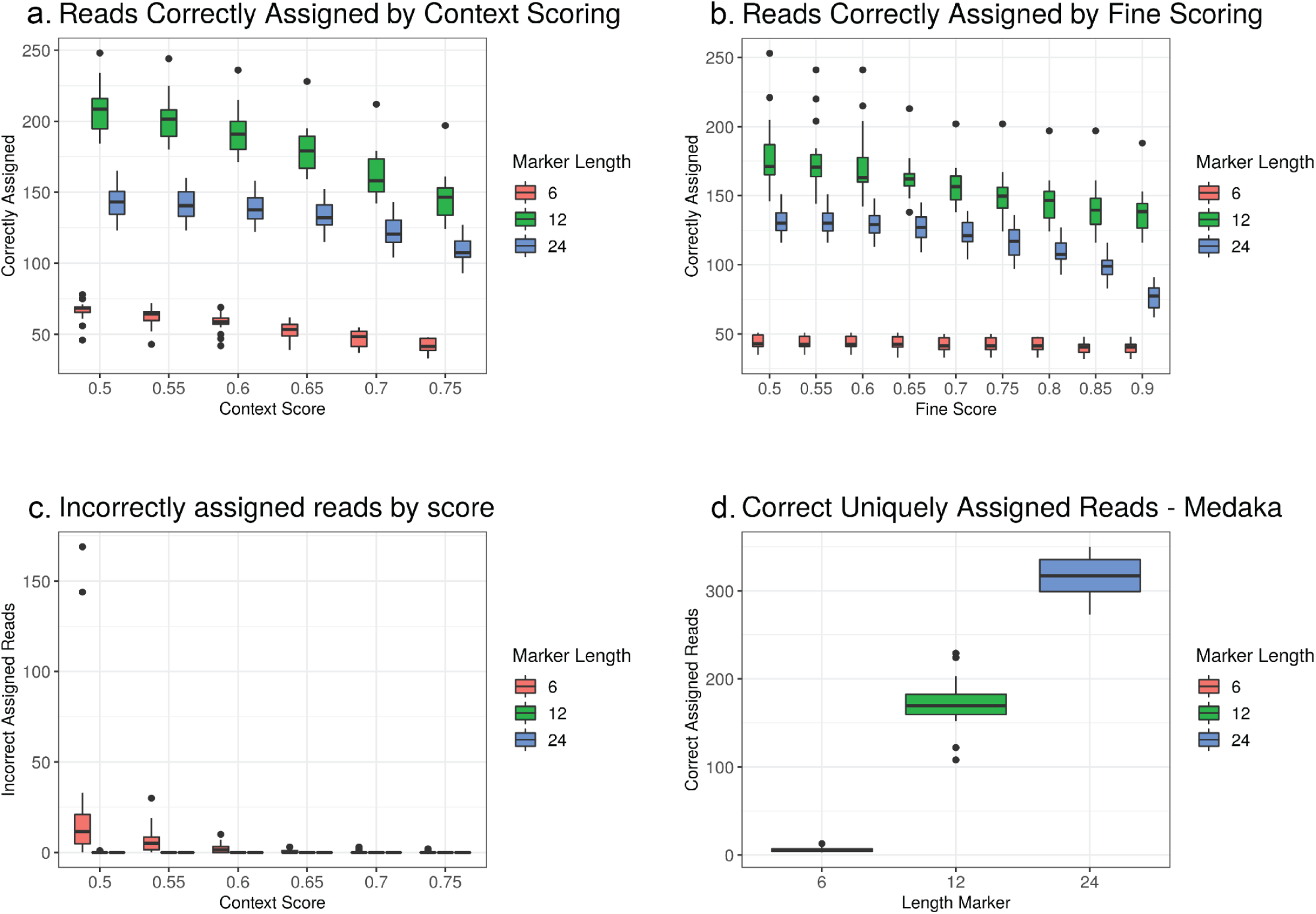
Number of correctly assigned reads for each of the 30 simulated plasmids containing a 6bp, 12bp, or 24bp unique region using different modes. Reads aligned using biobin mode (a,b) with a fixed fine score, comparing read counts at different context scores (a) or with a fixed context score, comparing read counts at different fine scores (b). The number of reads incorrectly assigned using different context scores (c). Count of uniquely mapping, correctly assigned reads using medaka mode (d).

**Supplemental Figure 3.**
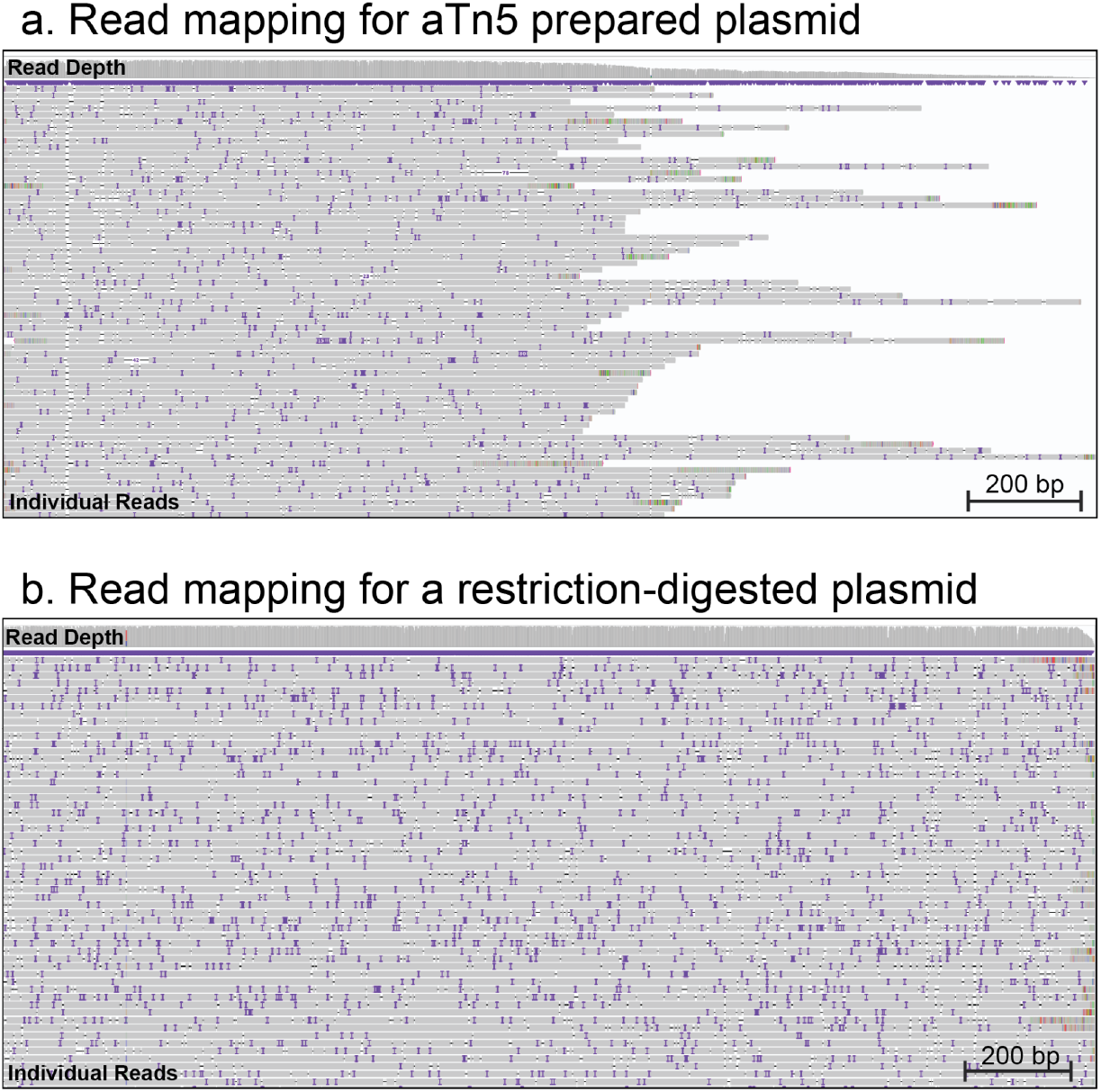
Difference in read coverage at plasmid ends with Tn5 vs restriction digest preparation. IGV view showing read coverage (uniquely mapped reads) for the end of a plasmid simulated to be prepared using Tn5 (a) or restriction digest (b) with read coverage indicated by height of gray panel at top.

